# Low-dose cadmium potentiates lung inflammatory response to 2009 pandemic H1N1 influenza virus in mice

**DOI:** 10.1101/346866

**Authors:** Joshua D. Chandler, Xin Hu, Eunju Ko, Soojin Park, Jolyn Fernandes, Young-Tae Lee, Michael L. Orr, Li Hao, M. Ryan Smith, David C. Neujahr, Karan Uppal, Sang-Moo Kang, Dean P. Jones, Young-Mi Go

## Abstract

**BACKGROUND:** Cadmium (Cd) is a toxic, pro-inflammatory metal ubiquitous in the diet that accumulates in body organs due to inefficient elimination. Many individuals exposed to dietary Cd are also infected by seasonal influenza virus. The H1N1 strain causes mild to severe pneumonia which can be fatal.

**OBJECTIVES:** To determine the influence of low-dose Cd exposure on inflammatory responses to H1N1 influenza A virus.

**METHODS:** We exposed mice to low-dose (1 mg CdCl2/l) Cd or vehicle (water) for 16 weeks prior to infection with a sub-lethal dose of H1N1. Lung inflammation was assessed by histopathology and flow cytometry. We used a combination of transcriptomics, metabolomics and bioinformatics to determine the molecular associations of inflammatory cells important in Cd-exacerbated responses.

**RESULTS:** Cd-treated mice had increased lung tissue inflammatory cells, including neutrophils, monocytes, T lymphocytes and dendritic cells, following H1N1 infection. Lung genetic responses to infection (increasing TNF-a, interferon and complement, and decreasing myogenesis) were also exacerbated. Global correlations with immune cell counts, leading edge gene transcripts and metabolites revealed that Cd increased correlation of myeloid immune cells with pro-inflammatory genes, particularly interferon-γ, and metabolites in amino acid, nucleobase, glycerophospholipid and vitamin B3 pathways.

**DISCUSSION:** Cd burden in mice increased inflammation in response to sub-lethal H1N1 challenge, which was coordinated by genetic and metabolic responses, and could provide new targets for intervention against lethal inflammatory pathology of clinical H1N1 infection.

## Introduction

Recent studies of the environmental toxic metal cadmium (Cd) show that levels found in human lungs potentiate pro-inflammatory signaling by the redox-sensitive transcription factor NF-κB (Chandler et al. 2016b; Olszowski et al. 2012). Cd is a commercially important metal used in NiCd batteries, plating, pigments and plastics (Kataranovski et al. 2009; Waalkes 2003; Zhang et al. 2014). Earlier occupational toxicities from Cd have been largely eliminated by occupational hygiene, and tobacco use remains as a major avoidable source of exposure. In nonsmokers, diet is the most important source of human Cd burden, on average representing about half of the exposure of individuals with active tobacco use (Riederer et al. 2013; Satarug and Moore 2004). Although Cd is poorly absorbed, there is no effective mechanism for elimination so that the biological half-life in humans is over 10 years (Satarug and Moore 2004; Suwazono et al. 2009). Our mechanistic studies show that low concentrations of Cd cause oxidation of thiol/disulfide redox systems (Go et al. 2011; Go et al. 2013a; Go et al. 2014), disrupt the actin-cytoskeleton regulation (Go et al. 2013a), cause translocation of thioredoxin-1 (Trx1) into the nucleus (Go et al. 2013a), increase NF-κB activity and increase pro-inflammatory cytokines (Go et al. 2011; Go et al. 2013b). Cd causes a dose-dependent oxidation of plasma cysteine/cystine (Cys/CySS) redox state (Eh CySS, Y.-M. Go, unpublished), the major low molecular weight thiol/disulfide system in plasma previously associated with pro-inflammatory signaling (Go and Jones 2005, 2011). Additionally, studies with a transgenic NLS-thioredoxin-1 (Trx1) mouse showed that increased nuclear Trx1 substantially potentiated adverse outcomes from H1N1 influenza infection (Go et al. 2011) and also increased oxidation of thiol/disulfide redox systems (Go et al. 2011).

H1N1 and other influenza A strains are highly infectious (Liu et al. 2015; Xing et al. 2015) and cause variable severity of disease (Baigent and McCauley 2003a, b; Safronetz et al. 2011a, b; Taubenberger and Kash 2010a, b). Host genetics impact severity of response (Horby et al. 2012, 2013), but inter-individual differences in sensitivity have mainly been linked to demographic factors, such as age and general health (Gagnon et al. 2013; Wilson et al. 2014). In most cases, the pandemic 2009 influenza A (H1N1) virus caused an uncomplicated respiratory tract illness with symptoms similar to those caused by seasonal influenza viruses. However, severe responses also occurred in a fraction of patients and were linked to increased levels of pro-inflammatory cytokines, termed a “cytokine storm” (Gautam 2012; Liu et al. 2016), as well as to excess activity of the complement system (Monsalvo et al. 2011). Cytokine storm included high levels of IL-6 and was strongly associated with decreased lung function and death (Bermejo-Martin et al. 2009; To et al. 2010).

The current study was designed to test whether Cd burden, at a level found in lungs of non-smokers, enhanced severity of HIN1 influenza virus infection in a mouse model. Mice were given water ad libitum with or without CdCl2 (1 mg/l) for 16 weeks, and 10 days before sacrifice each mouse was also intranasally exposed to either sub-lethal H1N1 or mock (sterile saline) treatment. Lung histopathology and immune cell counts were used to evaluate inflammatory responses and pathology. Gene expression analysis and high-resolution metabolomics were used to identify Cd-dependent associations of inflammatory cells using the open-source software, *xMWAS* (Uppal et al. 2018). Targeted analysis of IFNγ protein correlations with immune cells was used to confirm *xMWAS* results.

## Materials and Methods

### Animals

All experimental protocols for animal studies were approved by Emory University and Georgia State University Internal Care and Use Committees. Experimental methods were carried out in accordance with the relevant guidelines and regulations. Male C57BL6 mice (n = 11-13 per group) aged 8 weeks (Jackson Labs, Bar Harbor, ME, USA) were maintained in clean facilities and given sterile-filtered drinking water with 1 mg/l CdCl2 (Sigma-Aldrich, St. Louis, MO, USA) or vehicle for 16 weeks (including the 10-day duration of H1N1 infection). Food content of Cd (62 ± 1 ng/g food) was negligible compared to the Cd derived from water (Chandler et al. 2016b). Ten days prior to study completion, mice were inoculated with H1N1 (A/California/04/2009; dose of 0.6 x 10^3^ pfu) under isoflurane anesthesia. Day 10 was chosen as the time point of maximal sub-lethal morbidity but also as a clinically relevant time point for severe and persistent cases. People infected with influenza are generally asymptomatic for 1-4 days, then exhibit symptoms for 3-7 days (https://www.cdc.gov/flu/professionals/acip/clinical.htm). Virus titer from mouse lung homogenate is expressed as the log10 of the 50% egg infective dose (EID50) per ml homogenate.

For the main study, mice were weighed every 7 days until infection, then daily for the remaining 10 days. Mice were observed daily for other signs of influenza morbidity, e.g., a ruffled hair coat or labored breathing, and no apparent differences were observed between H1N1 and Cd+H1N1. Airway reactivity as measured by airway enhanced pause (PenH) was used to evaluate possible airway effects (Chandler et al. 2016a; Hwang et al. 2016). The PenH procedure does not require additional compound exposures (e.g., methacholine) which could impact the lung metabolome.

### Cd measurement by Inductively-Coupled Plasma Mass Spectrometry (ICP-MS)

Lung tissue ^114^Cd was assayed as previously described (Chandler et al. 2016b). ICP-MS procedures conformed to accuracy (100 ± 10%) and precision standards (relative standard deviation < 12%). Cd values were normalized to tissue mass. Human lung tissue samples were collected in the Emory Transplant Center (Emory IRB protocol 000006248) and maintained frozen at −80 °C until analysis.

### Flow cytometry and staining

Immune cells in lung tissue were quantified by fluorescence-activated cell sorting (FACS). Lung tissues were homogenized in DMEM and applied to Percoll gradients before washing with PBS. Total viable cells were counted using a hemocytometer with 0.4% trypan blue for dye exclusion. A subset of cells for FACS analysis was treated with anti-surface marker antibodies including CD11b (M1/70), CD45 (30-F11), B220 (RA3-6B2), Siglec F (E50-2440) (BD Pharmingen, Franklin Lakes, NJ, USA); CD11c (N418), F4/80 (BM8) (eBioscience, Thermo Fisher Scientific, Waltham, MA, USA); and CD103 (2E7), Ly6c (HK1.4), MHC-II (M5/114.15.2), CD3 (17A2), CD4 (GK1.5), and CD8 (53-5.8) (BioLegend, San Diego, CA, USA). Data were acquired using an LSR-II/Fortessa (Becton-Dickinson, Franklin Lakes, NJ, USA) and analyzed using FlowJo v. 10.1 (Tree Star, Inc., Ashland, OR, USA). CD11b^+^ DCs were gated as those with low CD103 expression, and vice-versa; we did not analyze a mixed population (CD11b^+^/CD103^+^). Absolute counts were expressed from the percentage from FACS analysis multiplied by viable cell counts obtained with a hemocytometer.

### Cytokine and chemokine assays

IFNγ was measured as previously described (Ko et al. 2016; Lee et al. 2015; Lee et al. 2016). Briefly, right lung lobe extracts were prepared by grinding and filter straining prior to IFNγ measurement according to manufacturer instructions (eBioscience, Thermo Fisher Scientific, Waltham, MA, USA).

### Analysis of glutathione and cysteine

Glutathione (GSH), oxidized glutathione (GSSG), cysteine (Cys) and oxidized form of Cys (cystine, CySS) in lung tissue and plasma were analyzed by HPLC with fluorescence detection (Go and Jones 2005; Jones et al. 2002; Jones and Liang 2009). Protein assay (bicinchonicic acid) was used to standardize the results to intracellular concentrations. Redox states of GSH/GSSG (Eh[GSSG]) and Cys/CySS (Eh[CySS]) were calculated using the Nernst equation (Go and Jones 2005; Jones et al. 2002; Jones and Liang 2009).

### Histopathology

The lung collected at 10 days after challenge was fixed with 10% normal buffered formalin (NBF) in PBS. After fixation NBF tissues were transferred to 70% alcohol and processed to paraffin using a Shandon Excelsior AS (Thermo Fisher Scientific). A standard program of dehydration was carried out in ethanol 70%, 80%, 90%, 95%, 100%, 100% 1 hour each, followed by xylenes twice for 30 min each. Finally, the tissue was soaked in paraffin for 30 min and repeated once, then held in the second paraffin bath for 1 hour. Left lung tissue sections (4 mice per group) were stained with hematoxylin and eosin (H&E) to assess histopathological changes as described (Ko et al. 2016). Images were acquired by Axiovert 100 (Zeiss, Oberkochen, Germany) at 100X magnification, and inflammation was blindly scored as previously described (Ko et al. 2016).

### Transcriptomics

20-25 mg of lung tissue was stored in RNAlater (Qiagen, Hilden, Germany) at −80 °C prior to lysis and RNA isolation using the miRNEasy kit (Qiagen, Hilden, Germany). 250 ng of RNA was prepared and hybridized to MoGene ST 2.0 arrays (Affymetrix, Thermo Fisher Scientific, Waltham, MA, USA). CEL files were converted to a Robust-Multi Array (RMA) and loaded in the gene set enrichment analysis (GSEA) Java applet (v. 2.2; Broad Institute) to test gene set enrichment. Annotations were made using MSigDB file ‘MoGene_2_0_st.chip’, accessed from the GSEA applet. Redundant annotations were collapsed using maximum intensities.

### High resolution metabolomics (HRM)

Lung tissue metabolites were extracted from 20-30 mg tissue in 15 μl/mg of 2:1 acetonitrile:water containing internal standards per mg tissue (Chandler et al. 2016a). Tissues were sonicated, vortexed and incubated on ice for 30 minutes before centrifugation at 16,000 g and 4 °C. Five μl of supernatant was injected on a Dionex Ultimate 3000 LC system coupled to QExactive High Field mass spectrometer (Thermo Fisher Scientific, Waltham, MA, USA); each analysis was performed with three technical replicates. Chromatographic separation was achieved with Accucore HILIC chromatography (100 x 2.1 mm, 2.6 μm, 80Å; Thermo Fisher Scientific, Waltham, MA, USA). Untargeted data were extracted with XCMS (Smith et al. 2006) and *xMSanalyzer* (Uppal et al. 2013), recovering 2,956 features (high resolution *m/z* peak paired with retention time). Feature intensities were log2-transformed prior to integrative network analysis using *xMWAS* (Uppal et al. 2018). The pathway enrichment analysis by *mummichog* software (Li et al. 2013) was used to identify metabolic pathways enriched in mass spectral features obtained from *xMWAS* analyses (Uppal et al. 2018); this approach was previously validated for relevant metabolites by coelution with pure standards and MS^2^ spectra (Frediani et al. 2014; Uppal et al. 2017).

### Data integration and network analysis

Immune cells (Cd+H1N1 significantly different from H1N1 alone), gene transcripts (from leading edges of gene sets significantly different between Cd+H1N1 and H1N1 alone) and lung *m/z* features (all) were selected for *xMWAS* analysis to identify Cd-dependent associations during H1N1 infection. *xMWAS* (Uppal et al. 2018) uses the sparse partial least squares (sPLS) regression method and the *network()* function implemented in R package mixOmics (Gonzalez et al. 2012; Le Cao et al. 2009) for generating the association matrices between datasets, with Student’s t-test used to evaluate the statistical significance of pairwise association scores.

### Statistics

Data are presented as the mean and standard error of the mean. One-way ANOVA with Holm-Sidak’s post-test was used to test statistical significance at a significance level of *p* < 0.05. The GSEA applet (v. 2.2; Broad Institute) produced all statistics for gene set enrichment (Subramanian et al. 2005). Genes were ranked according the “Signal2Noise” statistic with a significance threshold of false discovery rate (FDR)-adjusted *q* < 0.05. In *xMWAS,p* < 0.05 was the significance threshold for correlations.

### Data Sharing

In addition to Supplementary Figures, data corresponding to the Figures and Supplementary Figures have been provided as numbered Supplementary Files. Datasets corresponding to metabolomics and transcriptomics used for *xMWAS* analysis have also been included (see List of Supplementary Files).

## Results

### Selection of sub-lethal H1N1 dose

Selection of H1N1 virus titer for mouse infection was critical to achieve maximal sub-lethal morbidity as evaluated by body mass decrease and increased lung inflammation at Day 10 postinfection. We compared the effects of 5 different virus doses on severity of lung infection, measuring body weight change over 10 days and observing lung inflammatory histopathology at Day 10 (**Supplementary Figure 1A**). The results showed that the virus doses of 0.6-1.2 x 10^3^ pfu caused sustained body weight loss up to Day 10 post-infection. Lower doses exhibited body weight recovery after Day 8 or 9. Doses of 0.6-1.2 x 10^3^ pfu also caused extensive inflammatory peribronchiolar histopathology (**Supplementary Figure 1B**). We reasoned that the lowest dose that caused sustained body weight loss (0.6 x 10^3^ pfu) would enhance the ability to detect second-hit effects of Cd and selected this dose for the main study involving Cd pre-treatment before infection. Consistent with previous low-dose Cd studies (Thijssen et al. 2007), Cd pretreatment had no effect on body mass or weight gain compared to vehicle treatment in the weeks prior to infection (**Supplementary Figure 1C**).

### Impact of Cd treatment on pathology of H1N1 infection

Four groups of mice were maintained for the treatment period, half receiving vehicle (sterile drinking water) and half receiving low-dose Cd. At 10 days prior to termination, mice were intranasally exposed either to mock treatment with sterile saline (Ctl, Cd groups) or 0.6 x 10^3^ pfu of H1N1 (H1N1, Cd+H1N1). We examined lung histopathology to determine the extent of inflammation in airway (**Figure 1B**), blood vessels (**Figure 1C**) and interstitial spaces (**Figure 1D**). As expected, H1N1 infection caused inflammation to increase compared to control in all compartments (H1N1: airways, *p* = 0.0016; blood vessels, *p* = 0.0095; interstitial spaces, *p* = 0.0127; **Figure 1B-D**). Cd treatment alone did not significantly increase histopathological scoring, and combination of Cd treatment with H1N1 did not significantly increase histopathology score compared to H1N1 with vehicle treatment. However, the combination of Cd+H1N1 had consistently lower p-values compared to control than H1N1 alone, for each histological compartment (Cd+H1N1: airways, *p* = 2.6 x 10^-4^; blood vessels, *p* = 0.0011; interstitial spaces, *p* = 9.8 x 10^-4^; **Figure 1B-D**). Mice treated with Cd exhibited increased lung Cd burden compared to vehicle treatment (**Supplementary Figure 1D**). To compare our model to human lung Cd, we used the same analytical methods to measure Cd in available lung samples from both smokers (S) and non-smokers (NS) (**Supplementary Figure 1D**). Cd content (pmol Cd/g wet tissue, mean ± SE) in Cd-treated mouse lung tissue [Cd (n = 7), 241 ± 32; Cd+H1N1 (n = 7), 220 ± 29] overlapped the range of non-smokers (n = 5, 157 ± 29) and lower portion of the range for smokers (n = 14, 568 ± 137). Lung tissue Cd content in vehicle-treated mice was lower than in Cd-treated mice [Ctl (n = 7), 137 ± 70; H1N1 (n = 8), 24 ± 7]. Treatment with Cd had no effect on infection-mediated body weight loss (Day 10 percentage of Day 0 body weight at inoculation: H1N1, 78 ± 2%; Cd+H1N1, 79 ± 2%, **Supplementary Figure 1E**), infection-mediated airway enhanced paused (PenH; mean Day 10 results as percentage of Day 0 value: H1N1, 700 ± 85%; Cd+H1N1, 669 ± 56%, **Supplementary Figure 1F**) or lung virus titer (H1N1, 10.0 ± 0.3 log10[EID50/ml]; Cd+H1N1, 9.6 ± 0.4 log10[EID50/ml], **Supplementary Figure 1G**).

**Figure 1.**
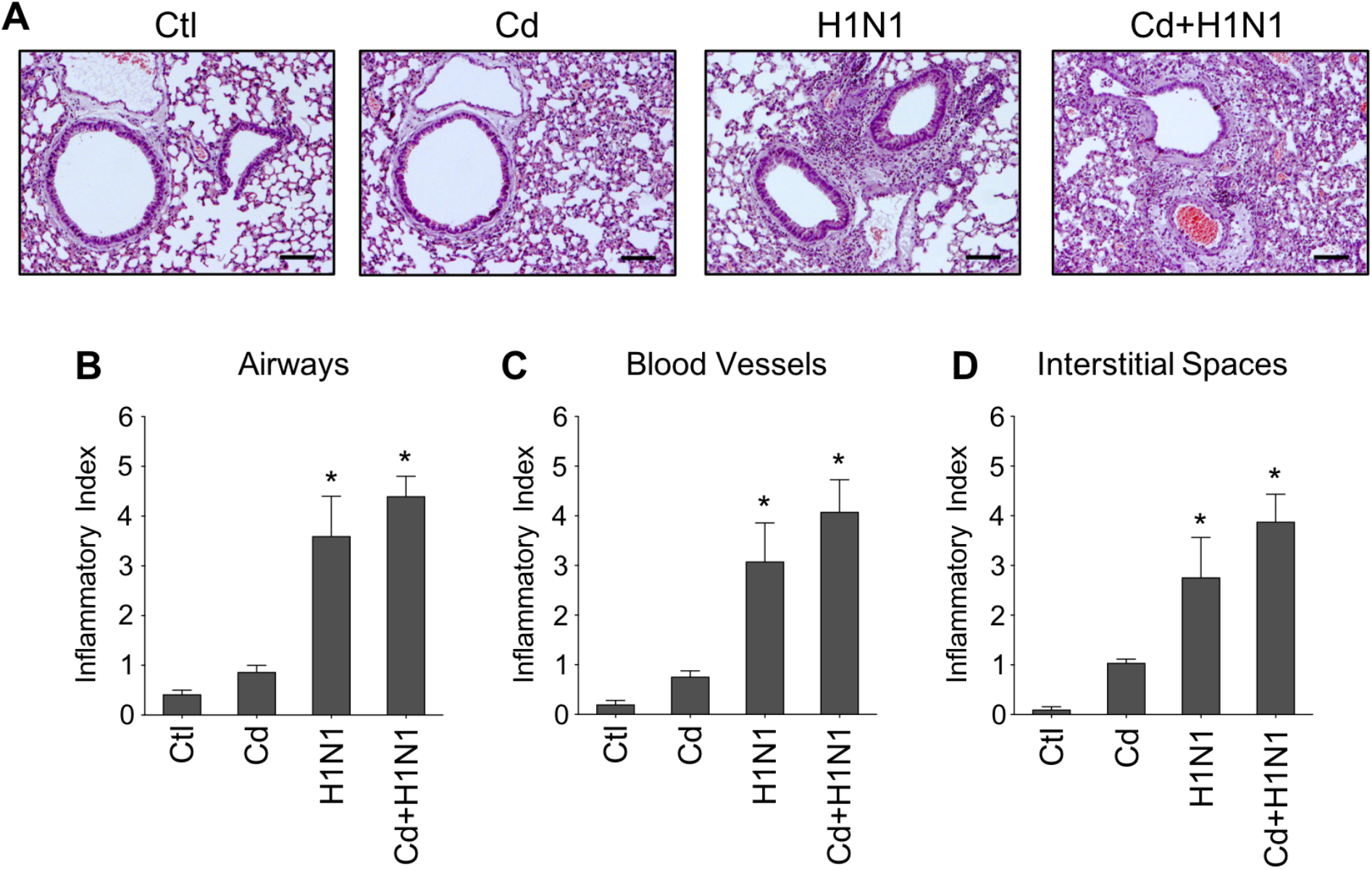
Impact of low-dose Cd on H1N1 infection lung histopathology. Mice were given ad libitum drinking water without or with CdCl2 (1 mg/l) for 16 weeks; ten days prior to completion, mice in both groups were given one intranasal dose of either sterile saline (Ctl, Cd) or H1N1 influenza virus (H1N1, Cd+H1N1). Lung tissue was collected and formalin-fixed upon sacrifice at Day 10 after infection. **(A)** Representative images of hematoxylin and eosin-stained sections of lung tissue are shown for each group (scale bar: 100 μm). **(B)** Airway, **(C)** blood vessel and **(D)** interstitial spaces were assessed for inflammatory foci in blinded fashion. n = 4-5 mice per group. \**p* < 0.05 compared to Ctl.

### Cd increased H1N1-mediated lung tissue inflammatory cells

Specific cellular contributions to lung inflammation were quantified using flow cytometry (**Figure 2**). Inflammatory immune cells were increased in lung tissue of Cd+H1N1 compared to H1N1, including neutrophils (*p* = 0.0002), monocytes (*p* = 0.0011), CD11b^+^ dendritic cells (DCs; *p* = 0.0001), CD103^+^ DCs (*p* = 3×10^-6^), plasmacytoid DCs (pDCs) (*p* = 7×10^-6^), CD4^+^ T lymphocytes (*p* = 0.0001) and CD8^+^ T lymphocytes (*p* = 0.0019) (**Figure 2A-G**). In contrast, we did not observe significant differences in the numbers of lung tissue alveolar macrophages or eosinophils when comparing Cd+H1N1 and H1N1 (**Supplementary Figure 2**).

**Figure 2.**
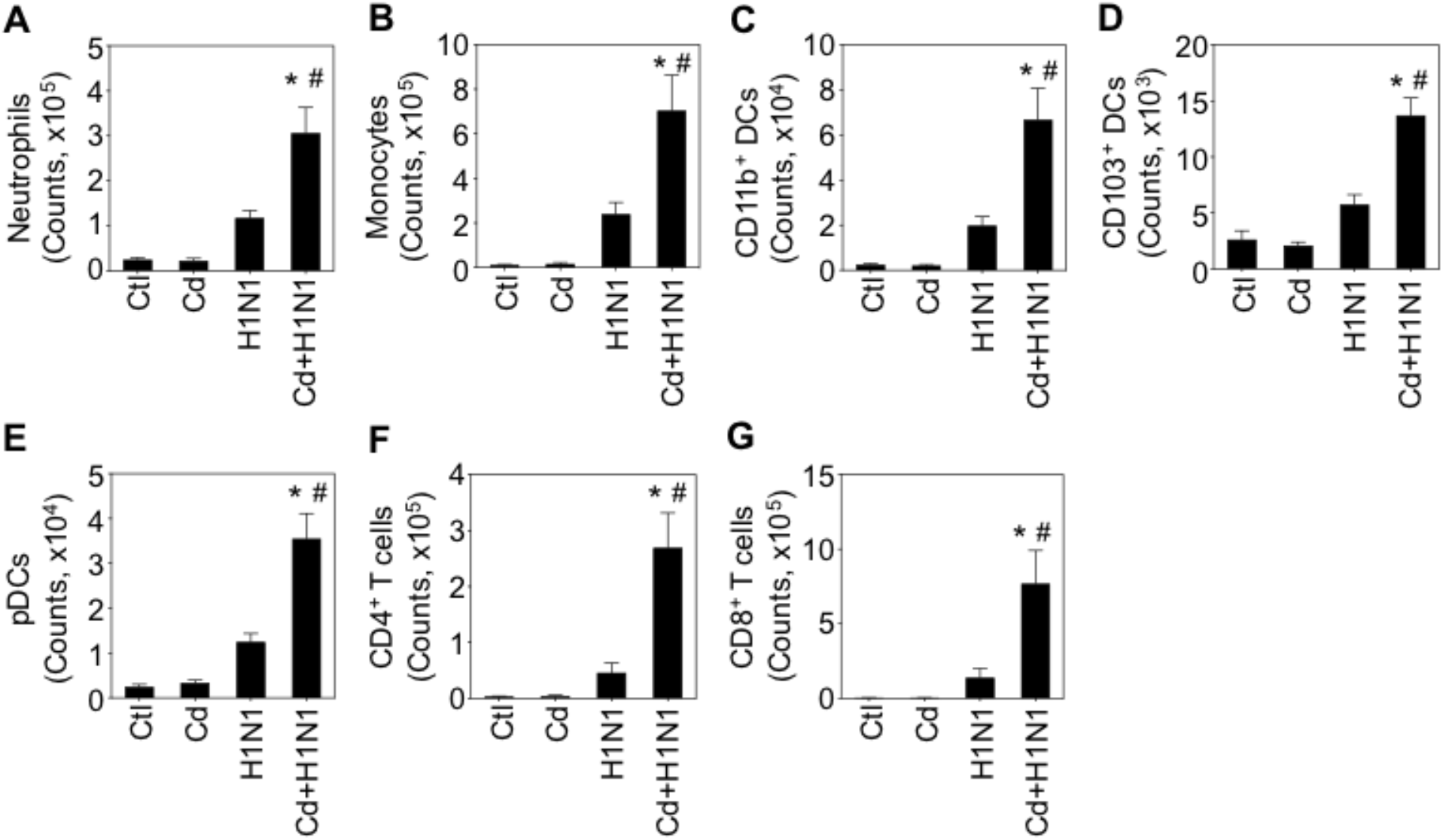
Cd increased lung immune cell response to H1N1 infection. Cells were isolated from lung tissue homogenates, labeled with appropriate antibodies for **(A)** neutrophils (PMNs), **(B)** monocytes, **(C)** CD11b^+^ dendritic cells (DCs), **(D)** CD103^+^ DCs, **(E)** plasmacytoid DCs (pDCs), **(F)** CD4^+^ T cells, (G) CD8^+^ T cells and quantified by flow cytometry. Bar plots show mean ± SEM of cell counts (n = 11-13, except for CD4^+^ and CD8^+^ T cells, n = 7-8). \**p* < 0.05 compared to Ctl, *#p* < 0.05 compared to H1N1 by one-way ANOVA with Holm-Sidak’s post-test.

### Impact of Cd on glutathione and cysteine in lung tissue and plasma

We measured the steady-state redox potentials of glutathione (Eh[GSSG]) and cysteine (Eh[CySS]) in plasma and lung tissue to establish whether low dose Cd treatment impacted these thiol-disulfide antioxidant systems in the context of sub-lethal influenza infection. H1N1 treatment did cause significant oxidation of Eh[CySS] in lung tissue (**Supplementary Figure 3B**), while no effects were observed for lung glutathione or for plasma (**Supplementary Figure 3A,C-D**). No potentiation of the effect noted in lung tissue Eh[CySS] by low dose Cd was observed.

### Cd enhanced pro-inflammatory gene set expression in H1N1-infected mice

To determine whether Cd impacted gene expression responses to H1N1 infection, we used Affymetrix MoGene ST 2.0 arrays followed by Gene Set Enrichment Analysis (GSEA) (Subramanian et al. 2005) to study effects on transcript abundances among the treatment groups, comparing overrepresentation of gene sets between experimental groups (n = 5). We selected GSEA gene sets with *p* < 0.05 and FDR *q* < 0.05 for gene set enrichment. Seven gene sets passed these criteria when comparing Cd+H1N1 and H1N1 (**Table 1**), including six that were overrepresented in Cd+H1N1 compared to H1N1 alone (1. allograft rejection, 2. complement signaling, 3. inflammatory response, 4. IFNa response, 5. IFNγresponse, 6. TNF-a signaling via NF-ĸB; each comprised predominantly of pro-inflammatory genes). The seventh gene set (myogenesis; comprised of cytoskeletal subunit genes) was overrepresented in H1N1 compared to Cd+H1N1. Each of the six gene sets overrepresented in Cd+H1N1 compared to H1N1 was also overrepresented in either of the H1N1-infected groups compared to mock-treated groups. In contrast, the myogenesis gene set was overrepresented in Ctl, Cd and H1N1 groups compared to Cd+H1N1.

**Table 1:** Cd increases pro-inflammatory gene enrichment in response to H1N1 challenge. Gene set normalized enrichment scores that are statistically significant comparing Cd+H1N1 and H1N1 (*p* < 0.05 and FDR *q* < 0.05) are shown. Normalized enrichment scores compared to controls (also *p* < 0.05 and FDR *q* < 0.05) are shown for H1N1 alone or Cd+H1N1. Negative enrichment values indicate *lower* gene abundances compared to the reference group (Ctl or H1N1, indicated in the column header). NA, gene set enrichment did not pass significance testing threshold.

| Gene Sets |  | Enrichment |  |  |
| --- | --- | --- | --- | --- |
| Name | Number of Genes | *H1N1* vs. *Ctl* | *Cd+H1N1* vs. *Ctl* | *Cd+H1N1* vs. *H1N1* |
| Allograft rejection | 189 | 3.02 | 3.21 | 2.31 |
| Complement | 181 | 2.36 | 2.56 | 1.73 |
| Inflammatory response | 199 | 2.61 | 2.75 | 1.84 |
| Interferon-α response | 89 | 2.79 | 2.88 | 2.04 |
| Interferon-γ response | 195 | 3.09 | 3.36 | 2.49 |
| TNF-α signaling via NF-κB | 196 | 2.60 | 2.82 | 2.06 |
| Myogenesis | 195 | NA | -1.66 | -2.14 |

The leading edge genes from each of the gene sets in **Table 1** were selected for global correlation analysis by *xMWAS* (Uppal et al. 2018). After accounting for redundancies across gene sets, this yielded 293 leading edge genes (**Supplementary File S3**). Two-hundred and twenty of these genes belonged to the pro-inflammatory gene sets in Table 1 that were increased in Cd+H1N1 compared to H1N1. Pro-inflammatory leading edge genes were predominantly comprised of cytokines, chemokines, receptors of cell activation and signaling proteins. The highest ranked leading edge gene from these sets (contributing most to their enrichment) was ‘interleukin 6 (interferon, beta 2)’ (*Il6*). The remaining 73 leading edge genes belonged to the myogenesis gene set with cytoskeletal subunit and regulatory genes including ‘actin, alpha 1’ (*Actal*), ‘creatine kinase, muscle’ (*Ckm)* and ‘collagen 1A1’ (*Collal*). The lowest ranked gene from this gene set (contributing most to the “negative enrichment” of the gene set in Cd+H1N1 compared to H1N1 alone; **Table 1**) was ‘myosin, light chain 7, regulatory’ (*Myl7*).

### Global correlation analysis of Cd effect on lung response to H1N1 influenza infection

To reveal organization of the network structure and pinpoint molecules critical to Cd-altered lung function, global correlations were made for immune cell counts, leading edge gene transcripts and metabolites using *xMWAS* (Uppal et al. 2018). *xMWAS* is a data-driven integration and network analysis tool which uses community detection to improve understanding of complex interactions. As input, we used the 293 leading edge genes from **Table 1** gene sets (**Supplementary File S3**) paired with inflammatory cells as measured in **Figure 2A-G** (**Suppelementary File S4**) and lung tissue metabolites measured by HRM (total of 2,956 metabolic features; **Supplementary File S5**) to produce experimental group-specific association networks, with the explicit goal to compare the respective associations of the H1N1 and Cd+H1N1 networks (**Figure 3**; n = 5). Student’s t-test was used to determine the association score (absolute value) threshold of 0.88 (determined by *xMWAS* software based on *p* < 0.05 criteria and n = 5 samples per group). The resulting network of H1N1 was comprised of 426 metabolic features, 160 gene transcripts and 7 immune cell types, while the network of Cd+H1N1 was comprised of 478 metabolic features, 158 gene transcripts and 6 immune cell types (**Figure 3A**; full network membership given in **Supplementary File S6**).

**Figure 3.**
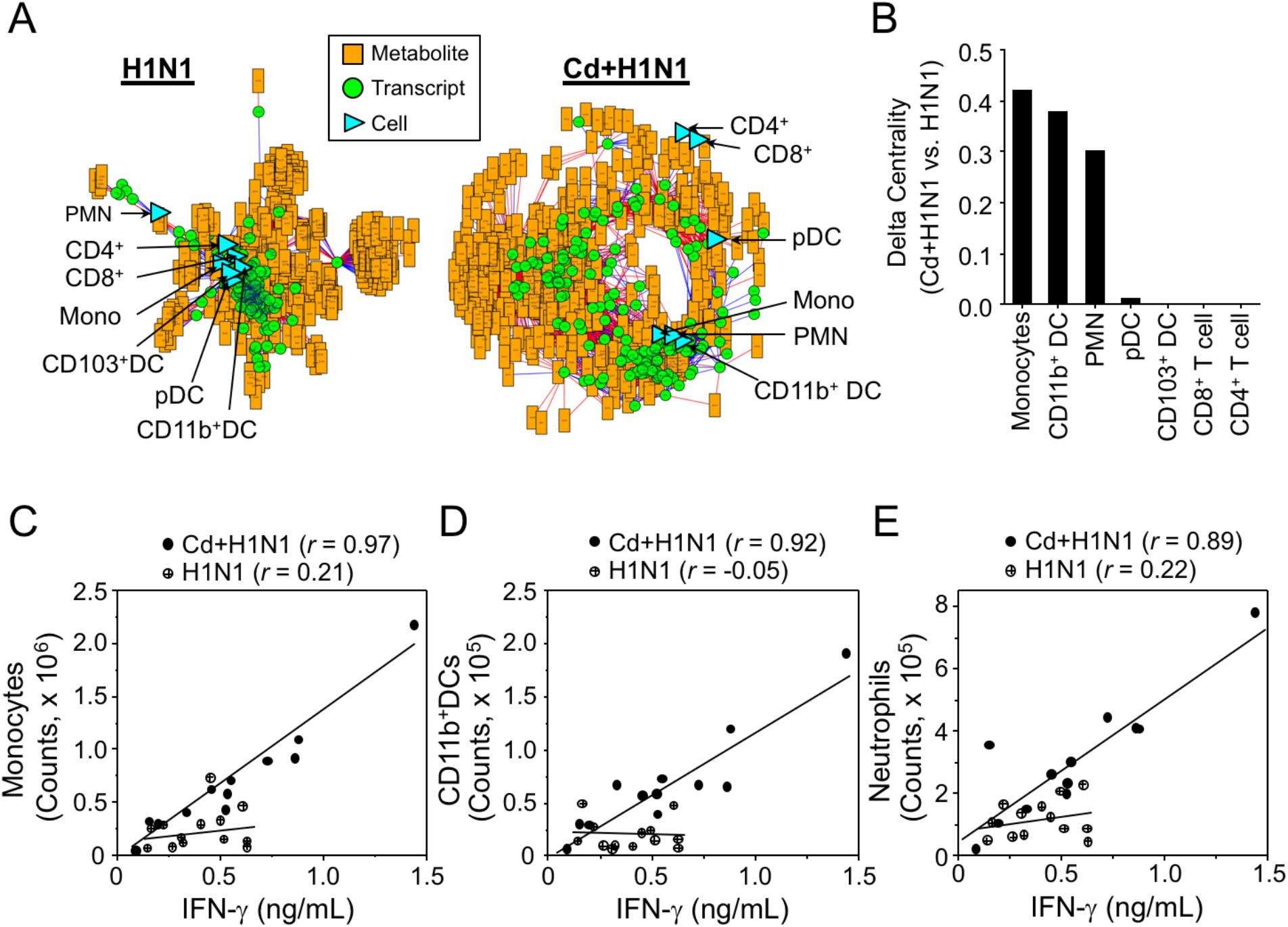
Cd promotes mouse lung myeloid cell responses to H1N1 infection. Inflammatory lung tissue cell counts (7 cell types, **Fig. 2**), leading edge gene intensities (293 leading edge genes selected by GSEA, **Supplementary File 3**;) and lung tissue metabolites measured by untargeted LC-MS (total of 2956 metabolic features, **Supplementary File 5**) were correlated using sparse PLS regression implemented by *xMWAS* (n = 5/group). **(A)** Networks were constructed of variables with inter-dataset absolute association score >0.88 at *p* < 0.05 (n = 5) for H1N1 or Cd+H1N1 groups. Symbols depict metabolic features (orange rectangle), gene transcripts (green circle) or cell counts (blue triangle), and lines indicate relationship between symbols (red, positive correlation; blue, negative correlation). The blue triangles have been enlarged for easier viewing. Abbreviations for cells are as follows: Monocytes, ‘Mono’; neutrophils, ‘PMN’; CD11b^+^ dendritic cells, ‘CD11b^+^ DC’; CD103^+^ dendritic cells, ‘CD103^+^ DC’; plasmacytoid dendritic cells, ‘pDC’; CD4 T lymphocytes, ‘CD4+’; and CD8 T lymphocytes, ‘CD8+’. Note, CD103^+^ DC is not found in the Cd+H1N1 network. **(B)** The absolute value of difference in eigenvector centrality (Delta Centrality) of each immune cell type was contrasted between the networks of Cd+H1N1 and H1N1 networks. Pearson’s correlation (*r)* was analyzed for the abundance of lung IFNγ protein and lung monocytes **(C)**, CD11b^+^ DCs (D) and neutrophils **(E)**. Open circles, H1N1 alone; closed circles, Cd+H1N1. The resulting statistics of correlation (r) and significance (*p*) were as follows: **C**, monocytes (Cd+H1N1, *r* = 0.97*,p* = 3×10^-7^; H1N1, *r* = 0.21, *p* = 0.49); **D**, CD11b^+^ DCs (Cd+H1N1, *r* = 0.92, *p* = 2×10^-5^; H1N1, *r* = −0.05, *p* = 0.86); **E**, neutrophils (Cd+H1N1, *r* = 0.89, *p* = 9×10^-5^; H1N1, *r* = 0.22, *p* = 0.48). n = 12-13.

To quantify the relative importance of “nodes” (cells, transcripts or metabolites) in each network, *xMWAS* calculates an eigenvector centrality score that can be compared across the groups (**Figure 3B**). Respective delta centrality scores showed that monocytes (+0.419), CD11b^+^ DCs (+0.380) and neutrophils (+0.301) had increased centrality in the Cd^+^H1N1 network compared to H1N1 (**Figure 3B**). Notably, these are all myeloid cell populations, so this result indicates an increase in myeloid cell importance in the Cd+H1N1 model. This result can also be qualitatively detected in the model visualizations, in which neutrophils, a major myeloid cell type, shift from peripheral (H1N1) to central (Cd+H1N1) network locations between the two experimental conditions, whereas CD4+ and CD8^+^ T lymphocytes, canonical lymphoid cells, do the opposite (**Figure 3A**). Additionally, CD103^+^, a lymphoid dendritic cell sub-population, was not found in the Cd+H1N1 network (indicating no strong correlations were made with this cell type in the network (**Figure 3A**). Thus, Cd treatment altered biological interactions of immune cells with gene transcript and metabolites in lung tissue following sub-lethal H1N1 infection, causing myeloid cell associations to be strengthened.

Previous research showed that influenza-induced IFNγ restricts protective innate lymphoid cell group II function in mouse lung following challenge with H1N1 (Califano et al. 2018). We therefore performed targeted tests to determine whether correlations with IFNγ protein, measured in lung tissue homogenate by ELISA, differed for myeloid and lymphoid cells in Cd+H1N1 compared to H1N1. Results showed that each myeloid cell count correlated with IFNγ only in Cd+H1N1 mice [monocytes (Cd+H1N1, *r* = 0.97, *p* = 3×10^-7^; H1N1, *r* = 0.21, *p* = 0.4862; **Figure 3C**), CD11b^+^ DCs (Cd+H1N1, *r* = 0.92, *p* = 2×10^-5^; H1N1, *r* = −0.05, p = 0.8657; **Figure 3D**) and neutrophils (Cd+H1N1, *r* = 0.89, p = 9×10^-5^; H1N1, *r* = 0.22, *p* = 0.4773; **Figure 3E**)]. Additionally, pDC exhibited a modest correlation with IFNγ in Cd+H1N1 not observed in H1N1 (Cd+H1N1, *r* = 0.69, *p* = 0.0122; H1N1, *r* = −0.12, *p* = 0.6869), while none of the lymphoid cell populations tested (CD8+, CD4+, CD103^+^ DCs) exhibited significant correlations with IFNγ in either H1N1 or Cd+H1N1 groups (**Supplementary Figure 4**). The results confirm the global correlation network analysis showing that Cd burden increases centrality of myeloid cells with pro-inflammatory pathways following H1N1 infection. Furthermore, the strength of the IFNγ correlations in Cd+H1N1 across cell types followed the same rank order as delta centrality (**Figure 3B**). These results implicate IFNγ signaling as highly relevant to increased inflammation and myeloid cell dominance caused by Cd treatment.

### Gene sets and metabolic pathways associated with Cd effect on immune cell responses to H1N1

We further examined the molecular composition of each of the correlation networks to determine which gene transcripts and metabolic pathways were central in each group. The overall network composition of Cd+H1N1 and H1N1 overlapped by >50% in both leading edge genes and metabolic features so we focused on the most divergent relationships between Cd+H1N1 and H1N1 by analyzing genes and metabolites with absolute delta eigenvector centrality, |Centralityed+H1N1 – CentralityH1N1|, ≥ 0.1. (If a gene or metabolic feature was not present in one of the correlation networks, its eigenvector centrality was considered 0.) Results for the transcriptome data showed 83 leading edge genes of ≥ 0.1 absolute delta centrality were present in the Cd+H1N1 and H1N1 networks (**Supplementary File S8**), with the IFNγ response gene set (27.4% of total gene counts; **Figure 4A**) being most represented. IFNa response (17.8%) followed, and myogenesis was least represented (5.2%). The IFNγ transcript, *Ifng*, had differential centrality of +0.435 for Cd+H1N1 network compared to H1N1. The transcript with most positive change for gene centrality (1.0) was immediate early response 2, *Ier2*, encoding a DNA-binding protein, and the transcript with most negative change for gene centrality (−0.88) was Dnaj (Hsp40) homolog, subfamily B, member 4, *DnajB4*, a chaperone protein targeted to the plasma membrane. Thus, the transcript data are consistent with Cd effects occurring through IFNγ-dependent signaling mechanisms and also involving more complex interactions of nuclear as well as plasma membrane systems.

**Figure 4.**
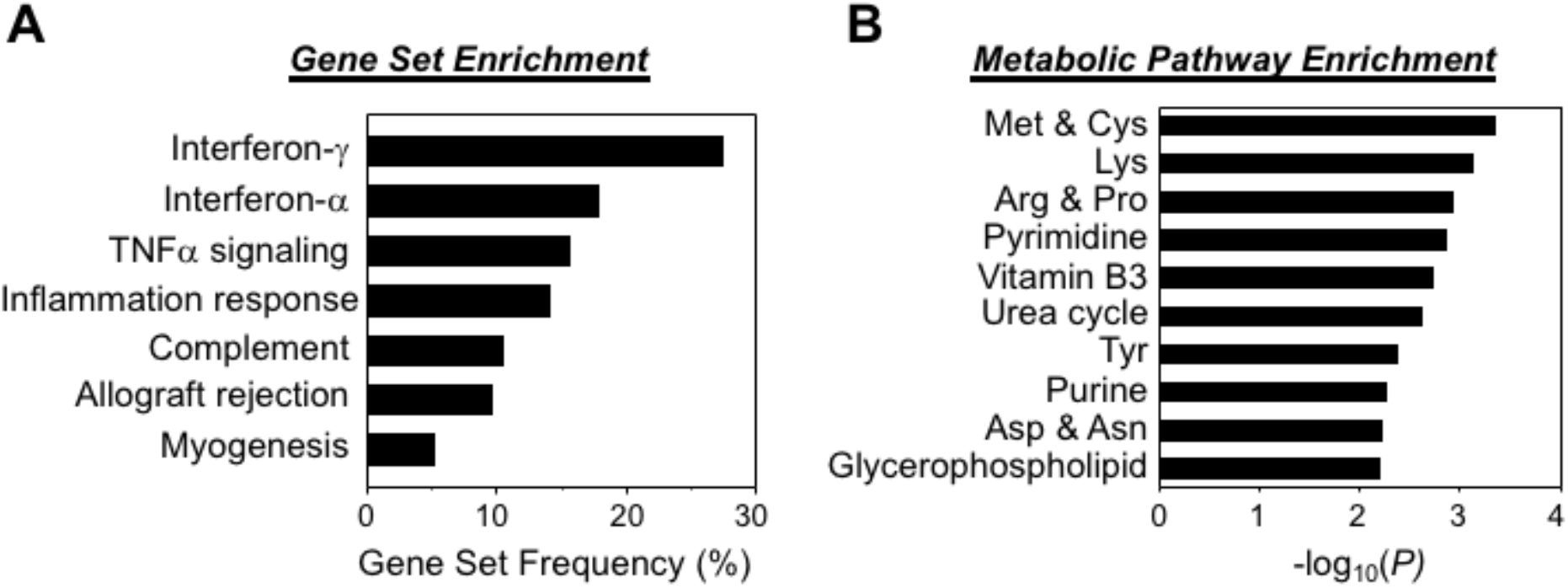
Cd potentiation of immune response to H1N1 infection through widespread effects of low Cd exposure on inflammation, mitochondrial function and regulation of immune cell populations through IFNγ. **(A)** Percentage of gene sets represented in the 83 leading edge genes (**Supplementary Table 2**) with delta eigenvector centrality (absolute value), |Centrality Cd+H1N1 – CentralityH1N1|, ≥ 0.1 between Cd+H1N1 and H1N1 networks. **(B)** Metabolic pathway enrichment analysis was performed on 165 metabolic features with delta eigenvector centrality ≥ 0.1 between networks of Cd+H1N1 and H1N1 using *mummichog.* The negative log10 of the pathway p-value is plotted as a bar graph for each significant pathway (*p* < 0.05, at least 3 metabolite matches in pathway).

Prior research has shown that low-dose Cd has widespread effects on metabolism, especially related to oxidative stress, mitochondrial β-oxidation of fatty acids and amino acid metabolism (Go et al. 2015). We analyzed 165 metabolic features with delta eigenvector centrality ≥ 0.1 (**Supplementary File S9**), using the metabolic pathway analysis software, *mummichog* 1.0.10 (Li et al. 2013). The top metabolic pathway was methionine and cysteine metabolism (**Figure 4B**). The largest positive centrality change for a single metabolite was for S-adenosylhomocysteine (+0.415), an intermediate in methylation processes linked to methionine. Other pathways included vitamin B3 (niacin/NAD), urea cycle and glycerophospholipid metabolism, all of which are connected to mitochondrial function and known to be perturbed by Cd. Other metabolic pathways included amino acid, pyrimidine and purine pathways that have previously been associated with inflammation and/or cell proliferation and turnover (**Figure 4B**). Consistent with this, the metabolite with the most negative centrality change was adenine (−0.953). The results are consistent with Cd potentiation of inflammation to H1N1 infection through widespread effects of low-dose Cd exposure on oxidative stress, mitochondrial function and regulation of immune cell populations through IFNγ.

## Discussion

A previous study showed that blood concentrations of toxic metals including Cd were significantly higher in H1N1 cases with pneumonia versus pneumonia not caused by H1N1, and two H1N1 cases having highest Cd concentrations did not recover from illness (Moya et al. 2013). Although limited by the small number of cases, the study suggested that Cd could be a risk factor for inflammatory response and enhanced severity of H1N1 virus infection-induced respiratory diseases. In the present study using a mouse model of H1N1 infection, we show that lung Cd burden, at a value present in human lung, exacerbated inflammation and lung injury from H1N1 infection in mice. Comparing Cd+H1N1 to H1N1-infected mice, Cd increased the Day 10 post-infection lung burden of myeloid (monocytes, neutrophils), lymphoid (CD4^+^ and CD8^+^ T cells) and multi-origin (DCs) bone marrow lineages of immune cells. Integrated network analysis of omics data further showed that Cd burden shifted the immune response from one centered on both lineages to one with myeloid cells (including CD11b^+^ DCs) as a dominant central hub. Thus, high Cd burden appears to contribute to worse outcomes following H1N1 infection by increasing the immune response with suppression of lymphoid and enhancement of myeloid cell participation in response to infection.

The global correlation network analysis of immune cells with gene transcript and metabolite abundances revealed the centrality of myeloid cells (monocytes, CD11b^+^ DCs [considered a myeloid DC subtype (Chistiakov et al. 2015)] and neutrophils) in the Cd+H1N1 group. Transcriptional responses suggest that Cd primes myeloid cells for tight coordination with IFNγ signaling. The finding that a unique lysosomal thiol reductase, IFNγ-inducible lysosomal thiol reductase, controls conditions for degradation of proteins to small peptides in lysosomes (Hastings and Cresswell 2011; Phan et al. 2002), raises the possibility that Cd specifically impacts IFNγ-dependent functions through interaction with, or oxidation of, the thioredoxin-like active site involved in disulfide reduction. Low dose Cd causes oxidation and disruption of the actin-cytoskeleton (Go et al. 2013a), suggesting that targeted effects on a critical signaling system could occur even without effects on major GSH or thioredoxin systems. Effects of low-dose Cd on the actin-cytoskeleton system triggers translocation of Trx1 to cell nuclei (Go et al. 2013a) and increase NF-κB activation (Go et al. 2011; Go et al. 2013b) and monocyte adhesion (Go and Jones 2005). Thus, these processes also could contribute to higher immune cell counts in Cd+H1N1 compared to H1N1. The GSEA results showing effects of Cd on pro-inflammatory gene sets (**Table 1**), as well as increased cytokine levels and immune cell counts, support such a Cd-dependent mechanism.

Alternative mechanisms for Cd effects on inflammatory pathology of H1N1 could involve cytoskeletal functions, myogenesis and elastogenesis in the inflamed and injured lungs. It is noteworthy that the combination of Cd pre-treatment with H1N1 infection was the experimental condition to cause a significant negative enrichment, or decrease in overall transcript levels, of the myogenesis gene set. Cytoskeletal regulation by tight junctions and adherens junctions is a key element of airway permeability to infiltrating immune cells (Rezaee and Georas 2014). We previously found that low dose Cd disruption of actin cytoskeleton regulation (Go et al. 2013a; Hu et al. 2017) triggered nuclear translocation of thioredoxin-1 and caused cell death (Go et al. 2013b). A study by Desouza *et al.* also showed that actin-binding proteins contribute to cell death, and actin dynamics plays a key role in regulating apoptosis signaling (Desouza et al. 2012). Studies of low dose Cd effects on actin cytoskeleton and cell death in fibroblasts, HeLa and HT-29 cells [(Go et al. 2013a; Hu et al. 2017); Y.-M. Go, D.P. Jones, unpublished] suggest that this could be a general cell response and contribute to barrier dysfunction. Thus, Cd could also contribute to severity of response through mechanisms involving impaired myogenesis, elastogenesis, increased cell death and failure to clear cell debris in the inflamed and injured lungs.

In contrast to the coherent picture provided by Cd effects on proinflammatory signaling, immune cell accumulation in the lungs and amplified effects on myeloid compared to lymphoid cells, metabolic effects associated with Cd effects were diverse and complex; the results suggested more of a generalized adaptive response than a causal role in exacerbation of inflammation. The top pathway from pathway enrichment analysis was for sulfur-containing amino acids which serve to support GSH synthesis and antioxidant activities. This would appear to be a protective mechanism, but maladaptive effects could also occur at specific subcellular sites. Pathway enrichment analysis also showed that pathways of several amino acids, nucleosides and phospholipids were overrepresented in the networks with differential eigenvector centrality ≥ 0.1 (**Figure 4B**). Overall, these results suggest that Cd burden causes a complex array of metabolic responses.

Inhaled Cd causes acute pneumonitis and lung edema (Nemery 1990), and intraperitoneal injection of Cd causes lung inflammation in rats (Kataranovski et al. 2009). However, chronic ingestion of Cd at levels mimicking those derived from the human diet has not previously been directly linked to inflammatory lung disease. The present research with 16 weeks of Cd in drinking water showed little evidence for inflammation without the second stimulus of H1N1 infection. Thus, Cd accumulation with advanced age (> 65 y), tobacco use, metabolic syndrome and/or obesity, may create a sub-clinical state poised to over-react to influenza virus infection-induced inflammation, which may in turn lead to worse morbidity or mortality (Choi et al. 2014; Fiore et al. 2011; Lee and Kim 2016; Olsson et al. 2002; Ramsey and Kumar 2013). For future research, the dose of virus used in our study permits recovery to study later complications such as fibrosis caused by low-dose Cd (Hu et al. 2017).

Finally, in light of common consumption of foods with relatively high Cd content (potatoes, cereals), the results indicate that studies in humans are needed to evaluate whether Cd burden is associated with severity of response to influenza infection. Cd burden can be readily evaluated by determination of Cd/creatinine level in urine (Haddam et al. 2011; Larsson and Wolk 2016). While there are no approved strategies to prevent Cd accumulation or decrease its burden, mechanisms discussed above could be targeted to prevent Cd-dependent potentiation of influenza severity. For instance, the mechanism for nuclear Trx1 translocation could be targeted to block excessive activation of transcriptional responses. Alternatively, Txnip, a vitamin D3-binding protein that controls nuclear Trx1 activity (Wang et al. 2002; Yamawaki et al. 2005), could provide a target for therapeutic manipulation. Trp14 catalyzes reduction of cystine (oxidized form of cysteine) (Pader et al. 2014) and could provide a target to protect against oxidation of proteins and downstream signaling and recruitment of immune cells. Cd also interacts with trace elements including selenium, zinc and iron (Gocmen et al. 2000; Riederer et al. 2013), providing additional strategies for supplement use (Choi et al. 2014; Moya et al. 2013).

In summary, this study showed that mice given Cd at a level achieved in humans from dietary intake had worse lung inflammation and inflammatory pathology after H1N1 influenza infection, in association with enhanced myeloid-dominant pro-inflammatory gene and metabolic responses. Studies are needed to evaluate whether Cd burden contributes to influenza severity in humans. Combined analyses of phenotypic measures with transcriptome and metabolome analyses enable global surveillance of complex systems responses to identify candidate mechanisms and potential therapeutic targets.

## List of Supplementary Figures and Data Files

**Supplementary Figure 1**. Virus dose response, response to Cd pre-treatment, and body mass, PenH and virus titer endpoints of main study.

**Supplementary Figure 2**. Flow cytometry-quantified eosinophils and alveolar macrophages.

**Supplementary Figure 3**. Steady-state redox potentials of glutathione and cysteine in lung tissue and plasma.

**Supplementary Figure 4**. Correlations of additional cell types with IFNγ.

**Supplementary File S1**. Corresponds to Figure 1 histopathology data.

**Supplementary File S2**. Corresponds to Figure 2 flow cytometry data.

**Supplementary File S3**. Leading edge genes selected for *xMWAS* analysis.

**Supplementary File S4**. Flow cytometry results selected for *xMWAS* analysis.

**Supplementary File S5**. Metabolomics dataset selected for *xMWAS* analysis.

**Supplementary File S6**. Corresponds to Figure 3 *xMWAS* analysis data and IFNγ correlations.

**Supplementary File S7**. Corresponds to Figure 4 enrichment tests data.

**Supplementary File S8**. Leading edge genes used for proportion testing following *xMWAS.*

**Supplementary File S9**. Metabolite features used for *mummichog* testing following *xMWAS.*

**Supplementary File S10**. Full Robust Multi-Array dataset from lung tissue Affymetrix array. Supplementary File S11. Corresponds to Figure S1 data.

**Supplementary File S12**. Corresponds to Figure S2 data.

**Supplementary File S13**. Corresponds to Figure S3 data.

**Supplementary File S14**. Corresponds to Figure S4 data.

## Acknowledgements

Dr. Young-Mi Go and Dr. Dean P. Jones share equal senior authorship in this collaborative research. We thank the Emory Integrated Genomics Core, Dana B. Barr PhD and Priya E. D’Souza MPH for technical assistance. This study was supported by NIEHS Grant R01 ES023485 (DPJ and YMG), R21 ES025632 (DPJ and YMG), NIH S10 OD018006 (DPJ), NIH/NIAID grants R01 AI105170 (SMK), R01 AI093772 (SMK), and R21 AI119366 (SMK), and NHLBI F32 1F32HL132493 (JDC) and Cystic Fibrosis Foundation CHANDL16F0 (JDC).

Author Contributions

- Designed research – YMG, DPJ, SMK
- Performed research – JDC, JF, XH, YTL, EJK, SJP, DCN, MRS, MLO, LH
- Contributed new reagents or analytic tools – KU
- Analyzed data – JDC, YTL, SJP, EJK, JF, XH, MRS
- Wrote the paper – JDC, YMG, DPJ

## Abbreviations

Cd: Cadmium;
Ctl: control;
Cys: cysteine;
CySS: cystine;
FACS: fluorescence-activated cell sorting;
FDR: False discovery rate;
GSEA: Gene set enrichment analysis;
H1N1: pandemic 2009 influenza A (H1N1) virus;
HRM: High resolution metabolomics;
ICP-MS: Inductively coupled plasma mass spectrometer;
IFNγ: interferon gamma
IL-1β: interleukin-1beta;
LC: liquid chromatograph;
MS: mass spectrometer;
*m/z*: mass to charge;
RT: retention time;
sPLS: sparse partial least squares;
Trx: thioredoxin

